# Transcriptome Analysis of Bael (*Aegle marmelos* L.) a Member of Family Rutaceae

**DOI:** 10.1101/346403

**Authors:** Prashant Kaushik, Shashi Kumar

## Abstract

*Aegle marmelos* is a medicinally and horticulturally important tree member of the family Rutaceae. It is native to India where it is also known as Bael. Despite its importance; the genomic resources of this plant are scarce. This study presented the first-ever report of expressed transcripts in the leaves of *Aegle marmelos*. A total of 133,616 contigs were assembled to 46,335 unigenes with the minimum and maximum lengths of 201 and 14,853 bp. There were 7002 transcription factors and 94,479 simple sequence repeat (SSR) markers. The *A. marmelos* transcripts were also annotated based on information from other members of Rutaceae; namely *Citrus clementine* and *Citrus sinensis*. A total of 482 transcripts were annotated as cytochrome p450s (CYPs) and 314 transcripts were annotated as glucosyltransferases (GTs). In the *A. marmelos* leaves the monoterpenoid biosynthesis pathway was predominant. This study provides an important genomic resource along with useful information about *A. marmelos*.

## 1. Introduction

*Aegle marmelos* (2n = 18) or Bael is an underexploited member of family Rutaceae. Believed to be native to the Indian subcontinent, it is well distributed throughout the tropical and subtropical belts of southeast Asia [1,2]. Botanically, *A. marmelos* is a deciduous tree stretching up to 10 m in height that flowers during the months of May – June [3,4]. It is also commonly grown as a horticultural plant in India and its fruits are processed as juice or candies, as well as eaten fresh. During the past few decades, a spike in cultivation as a horticulture plant has been attributed to it has medicinal properties, along with a hardy nature allows it to be cultivated on marginal lands with acidic or alkaline soils [5,6].

The traditional medicine system of Ayurveda in India routinely uses every part of *A. marmelos* as a therapy of medical conditions [7,8]. The leaves are most easily accessible and therefore regularly used for the treatments than any other plant part. *A. maremelos* leaves are used to treat jaundice and help in wound healing when applied as a paste on a wound surface [9]. Moreover, *A. marmelos* leaf extracts have been proved to be a better cure for gastrointestinal and hematopoietic damage than its fruits [10]. Leaf extract of *A. marmelos* is used as a medication against a number of chronic diseases like diabetes, pancreatic cancer, and arthritis [11–14]. All these medicinal properties of *A. marmelos* leaves are attributed to various phytochemicals present in the leaves such as aeglin, rutin, γ-sitosterole, β-sitosterol, euginol, marmesinin, glycoside, skimmianine, etc. Broadly these phytochemicals can be divided into three main classes viz., alkaloids, phenylpropanoids and terpenoid [15,16].Furthermore, *A. marmelos* leaf extract is used for the green synthesis of gold and silver nanoparticles [17,18].

The sum-total of all the transcripts captured in the cell of an individual organism is called its transcriptome [19]. There are two ways to capture the expressed transcripts: either by microarray, which is limited to predefined sequences, or by performing RNA-Seq using second generation sequencing technologies [20]. This kind of sequencing has revolutionized the understanding of non-model organisms and has evolved as one of the first choices of methods to apply to gene discovery and expression profiling of non-model organisms [21,22]. The availability of well-defined computational tools along with well-applied methodology has further demonstrated the effectiveness of *de novo* transcriptome assemblies in organisms even without a reference genome [23,24].

Genomic resources in *A. marmelos* are scarce as compared to other members of Rutaceae, like *Citrus sinensis* (Sweet Orange) and *Citrus clementina* (Clementine), both having well-annotated genomes [25,26]. To the best of our knowledge, this is the first detailed report on the transcriptome of this medically important plant. Moreover, only 6 ESTs are available in the National Center for Biotechnology Information (NCBI) database (Accessed on 25 May, 2018). An investigation into the leaf transcriptome of *A. marmelos* can help to answer key questions regarding various aspects related to genes and their gene function, via the pathways involved in the metabolic compound formation. Therefore, we used RNA sequencing followed by *de novo* transcriptome assembly of *A. marmelos* leaves to identify the transcription factors, simple sequence repeats (SSRs), and transcripts related to important metabolic pathways in the leaves of *A. marmelos*. Also, the information regarding cytochrome P450s (CYPs) and Glucosyltransferases (GTs) extant in the leaf of *A. marmelos* was also accomplished.

## 2. Materials and Methods

### 2.1. RNA Isolation and Sequencing

Yung and tender leaves from three mature and healthy plants of *A. marmelos* variety “Kaghzi” (~ 5 years old) were collected from the Government Garden Nursery (coordinates at 29°58′06.9”N 76°52′50.8”E) in Haryana, India. The sampled leaf tissues were stored in RNAlater (Life Technologies, USA) till further use. RNA was extracted with a TRIZOL-based RNA extraction protocol for plant leaves [27,28]. The quality of the extracted RNA was checked on a 1% formaldehyde denaturing agarose gel and further quantified using a Nanodrop ND-1000 spectrophotometer (Nanodrop Technologies, Montchain, DE, USA). A pooled sample of RNA from three selected plants was used for a single cDNA library preparation. The library was prepared with a TruSeq RNA Library Prep Kit v2 from Illumina^®^ (Illumina, Inc., USA) and the library quantification was done using a Qubit Fluorometer (Qubit™ dsDNA HS Assay Kit) and Agilent D1000 ScreenTape system (Agilent Technologies, Santa Clara, CA, USA). The library was further sequenced on the Illumina HiSeq 2500 (2×150 bp) platform (Illumina, Dedham, MA, USA).

### 2.2. De Novo Assembly and Identification Coding Sequences

The cleaned reads were assembled using Trinity software (version 2.4.0) [29] and TransDecoder v. 3.0.1(http://transdecoder.sourceforge.net/) was used to identify candidate coding regions within the generated transcripts and to look for open reading frames (ORF) of at least 100 amino acids long to decrease the chances of false positives.

### 2.3. Gene Function Annotation

The transcripts with ORFs were annotated with BLASTX (default parameters, e-value cut-off 10^-5^) by resemblance counter to NR (NCBI non-redundant protein sequences database of NCBI), protein family (Pfam), Kyoto Encyclopaedia of Genes and Genomes (KEGG) (http://www.genome.jp/kegg/), e-value cut-off of 1 × 10^−5^ and Cluster of Orthologous Groups COG (https://www.ncbi.nlm.nih.gov/COG/) [30]. The gVolantes server (https://gvolante.riken.jp) was used for the assessment of the completeness of transcriptome assembly via BUSCO_v3 selecting the plant ortholog set. Only transcripts pertaining to plant species were extracted and used for gene ontology. Pfam annotation was done with Hmmerscan while Blast2Go was used for Gene Ontology (GO) annotation [31,32]. KEGG orthologies were estimated using KEGG Automated Annotation Server (KAAS) by means of single-directional best hit method (http://www.genome.jp/kegg/kaas/).

### 2.4. Identification of Transcription Factors

Transcription factors families present in the leaves of *A. marmelos* were identified by searching coding sequences identified by Trans Decoder against the plant transcription factor database (PlnTFDB) (http://plntfdb.bio.uni-potsdam.de/v3.0/downloads.php) with an e-value cut-off of < 1 e^−10^.

### 2.5. Identification of simple sequence repeats (SSRs)

The presence of SSRs was determined by using MIcroSAtelliteidentification tool v1.0 (MISA) (http://pgrc.ipk-gatersleben.de/misa/). Briefly, the transcripts were checked 10 times for mono-repeats, 6 times for di-repeats and 5 times for tri/tetra/penta/hexa-repeats.

## 3. Results

### 3.1. De novo Assembly, Gene Prediction, and Functional Annotation

RNA-Seq targeting expressed coding sequences has been used successfully in many medicinal and non-model plant species that do not have a reference genome (e.g., *Prosopis cineraria* [33], *Andrographis paniculate* [34], *Phyllanthus emblica* [35], *Picrorhiza kurroa* [36], *Azadirachta indica* [37]). Moreover, being a tree, *A. marmelos* can have a large genome size which further restricts genome sequencing efforts [38].

The pooled RNA sample of *A. marmelos* leaves of RIN value around 8.0 generated a total of 115.92 million paired reads of high-quality (Phred score>30). Trinity assembler was used for the assembly and after trimming of adapters a total of 133,616 contigs (only from reads of 200 bp and above in length) clustered into 46,345 unigenes (Table 1). The raw data obtained as a result of sequencing was submitted to NCBI BioProject (PRJNA433585). The assembly completeness report from gVolante estimated that the transcriptome assembly was 90.15 percent complete. We scrutinized for an open reading frame of at least 100 amino acids long to decrease the chances of false positives during open reading frames (ORF) predictions. The annotated transcripts with ORFs are listed in Table 2. A total of 90,525 transcripts were annotated to GO terms. The transcripts related to plant species were extracted and used for gene ontology (Table 2).

**Table 1.**
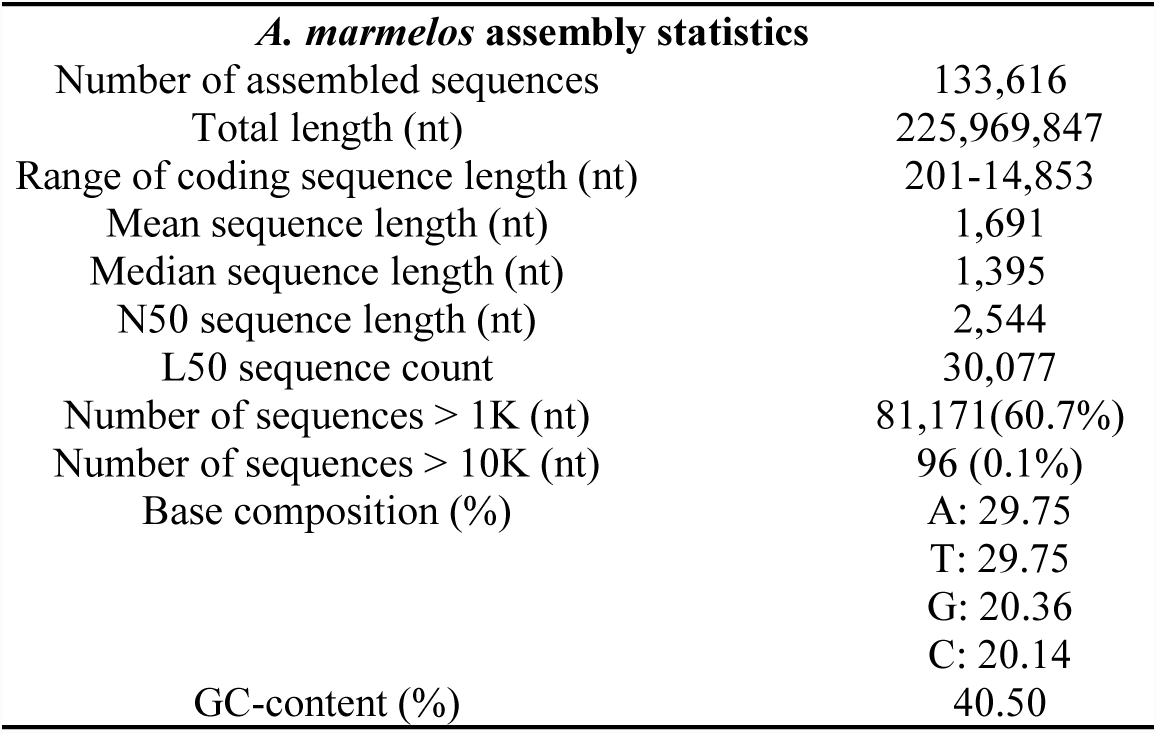
Assembly statistics of the leaf transcriptome.

| *A. marmelos* assembly statistics |  |
| --- | --- |
| Number of assembled sequences | 133,616 |
| Total length (nt) | 225,969,847 |
| Range of coding sequence length (nt) | 201-14,853 |
| Mean sequence length (nt) | 1,691 |
| Median sequence length (nt) | 1,395 |
| N50 sequence length (nt) | 2,544 |
| L50 sequence count | 30,077 |
| Number of sequences > 1K (nt) | 81,171(60.7%) |
| Number of sequences > 10K (nt) | 96 (0.1%) |
| Base composition (%) | A: 29.75 |
|  | T: 29.75 |
|  | G: 20.36 |
|  | C: 20.14 |
| GC-content (%) | 40.50 |

**Table 2.** Annotation summary of *A. marmelos* leaf transcripts.

| Parameters | Numbers |
| --- | --- |
| Total Transcripts | 133,616 |
| Total ORFs | 165,230 |
| Transcript BLASTX (Plant Species) | 126,101 |
| Transcript annotated with GO terms | 90,525 |
| Transcript BLASTP | 82,445 |
| Transcript BLASTX against Pfam | 67,451 |
| Transcript annotations against KEGG | 83,221 |
| Transcript annotated with COG | 81,636 |

### 3.2. GO Annotation

In total 600,642 Gene Ontology (GO) terms were mapped to the *A. marmelos* leaf contigs belonging to all the three possible classes, i.e., biological process (227,921 transcripts), cellular component (188,465 transcripts) and molecular function (184,25 transcripts) in the GO database (Figure 1). The breakdown of the proteins associated with the various biological process, cellular components and molecular functions is illustrated in Figure. 2. The “integral component of membrane” (GO: 0016021) associated with various cellular components, “transcription DNA-templated” (GO: 00006351) associated with biological processes and “ATP binding” (GO: 00005524) associated with molecular function were the most mapped terms in their respective categories (Figure. 2).

**Figure 1.**
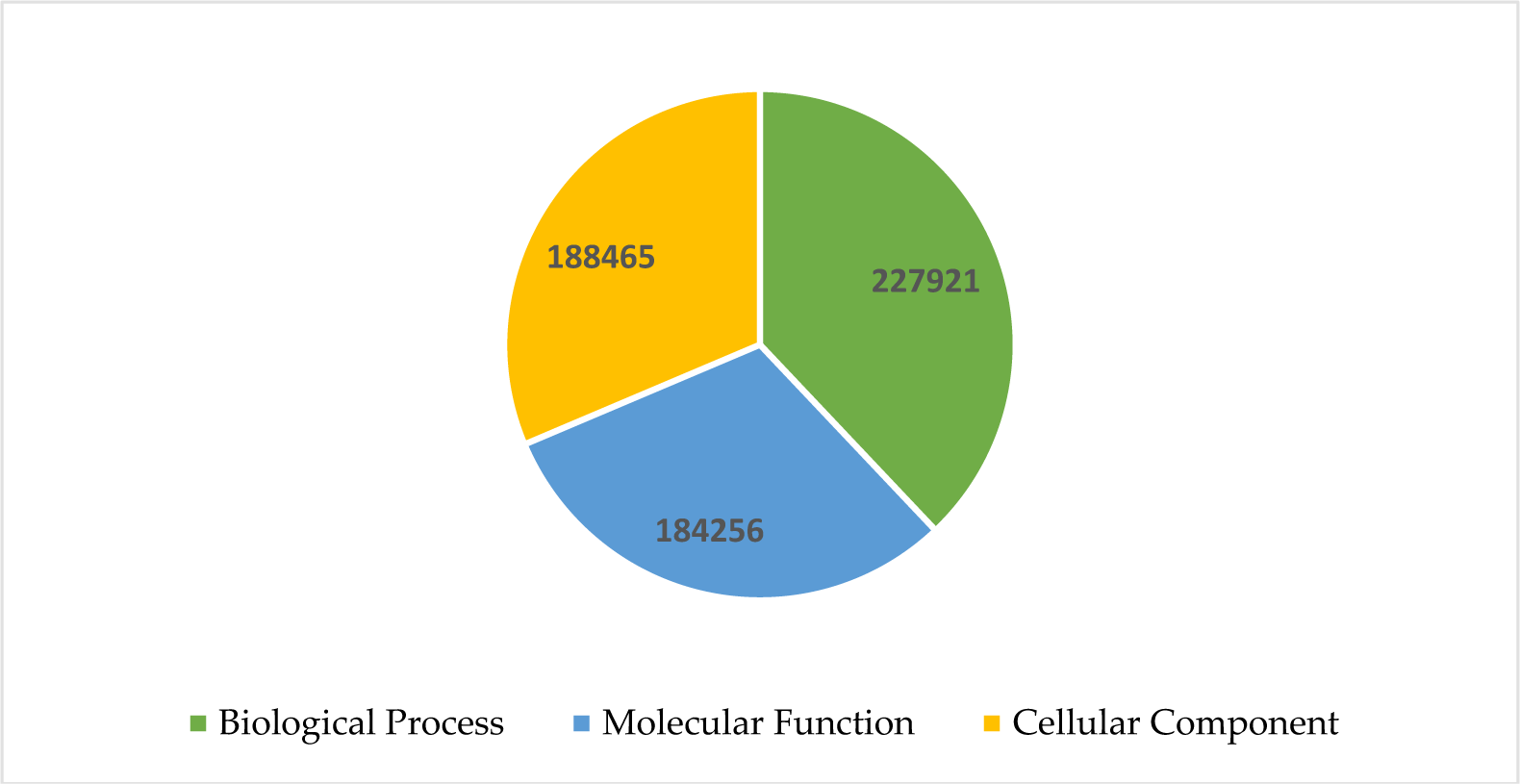
Genes associated with the biological process, cellular components, and molecular functions in the *A. marmelos* leaf transcriptome assembly.

**Figure 2.**
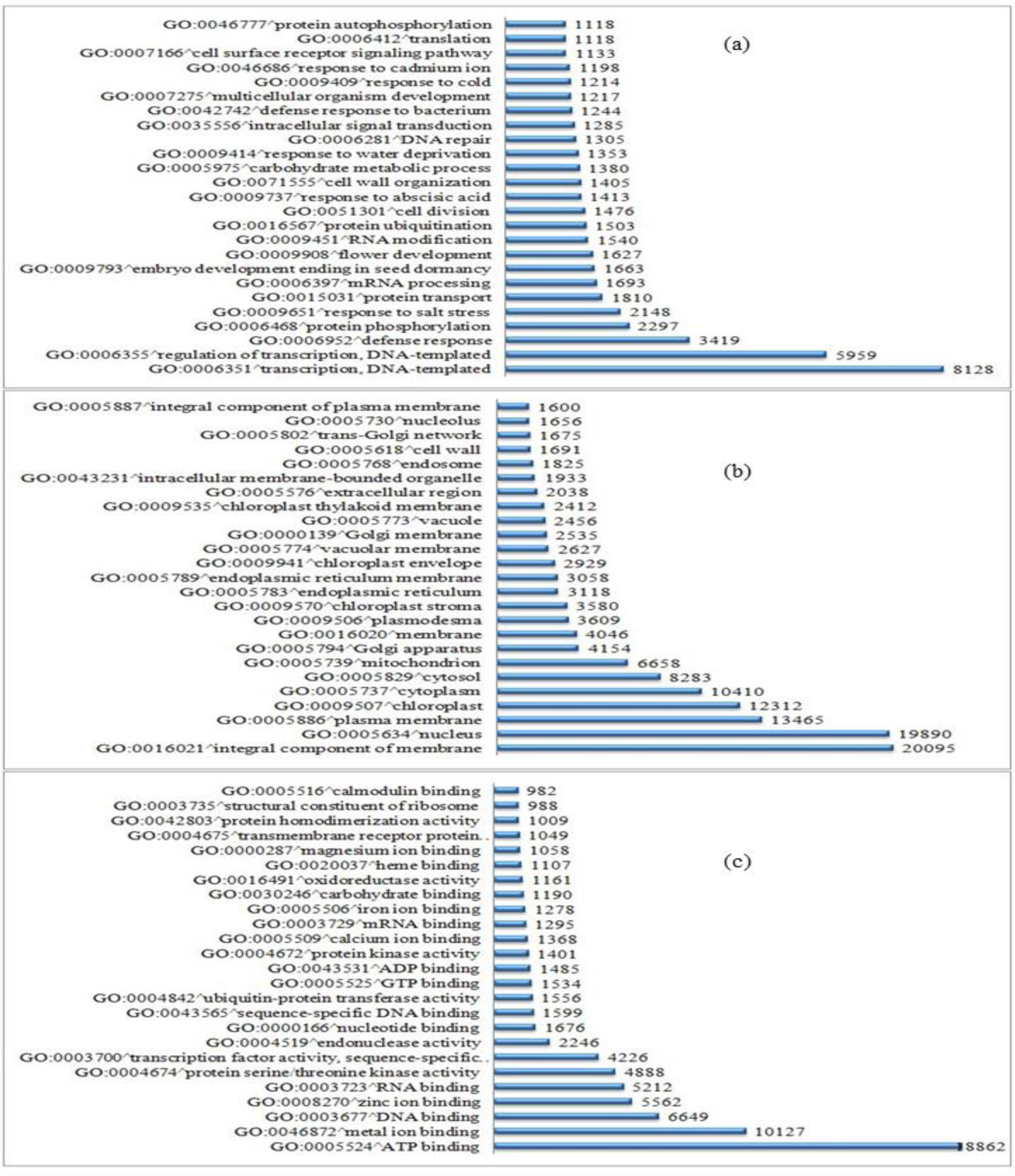
Gene Ontology (GO) classification of *A. marmelos* transcripts. GO term are divided in three main categories; biological process(a), cellular component (b), and molecular function (c).

The GO terms primarily define three categories of functions namely biological, cellular, and molecular functions for a gene product. This is achieved by associating a gene with their ontologies [39,40]. Earlier studies have pointed out a higher metabolic activity in the leaves of *A. marmelos* which is because of the presence of phytochemicals like alkaloids, flavonoids and phenols [41,42]. We have identified a number of GO terms in the leaves of *A. marmelos* this information could lead to the identification of important pathways of metabolic compounds in *A. marmelos* [43].

### 3.3. Citrus database annotation

The *A. marmelos* transcripts were also annotated via Phytozome (https://phytozome.jgi.doe.gov/) with reference to *Citrus clementina* and *Citrus sinensis* genome. This resulted in the mapping of 78.44 % of the transcripts to the *Citrus clementine* and 79.85 % to the *Citrus sinensis* genome (Table 3). An almost similar number of transcripts were annotated with GO terms and KEGG annotation respectively. Though, recently an extensive amount of relatedness was observed within the members of genus Citrus of family Rutaceae based on the study performed by using the whole genome sequences of 60 members in the Citrus genus, the authors even pointed out the need for reformulation of the genus Citrus this could also rationalize our almost similar annotation results of *A. marmelos* transcripts with *Citrus clementina* and *Citrus sinensis* genomes [44].

**Table 3.**
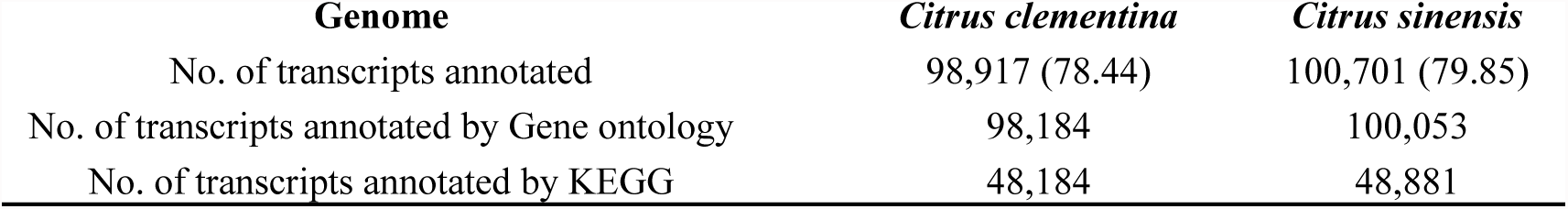
Annotation summery of *A. marmelos* leaf transcripts with *Citrus sinensis* and *Citrus clementina* genome.

| Genome | *Citrus clementina* | *Citrus sinensis* |
| --- | --- | --- |
| No. of transcripts annotated | 98,917 (78.44) | 100,701 (79.85) |
| No. of transcripts annotated by Gene ontology | 98,184 | 100,053 |
| No. of transcripts annotated by KEGG | 48,184 | 48,881 |

### 3.4. Simple Sequence Repeats (SSRs) Prediction

Simple sequence repeats (SSRs), or short tandem repeats or microsatellites, are short repeat motifs that show length polymorphism due to insertion or deletion mutations of one or more repeat types [45]. We analyzed for the abundance of SSRs of annotated plant transcripts for *A. marmelos* leaf transcripts using the MISA tool and the predicted SSRs statistics are shown in Figure 3. There were 58,354 transcripts which contained SSRs and among these, 23,034 had more than one SSRs. In total 94,479 SSRs were identified of which 65.27 % were mono repeats, 19.78 % were di repeats, and 13.40 % were tri repeats (Figure 3). Tetra, penta and hexa repeats were 1.01, 0.26 and 0.24 %, respectively (Figure 3). However, out of total 94,479 SSRs identified 11,400 (12.06 %) were related to the compound formation.

**Figure 3.**
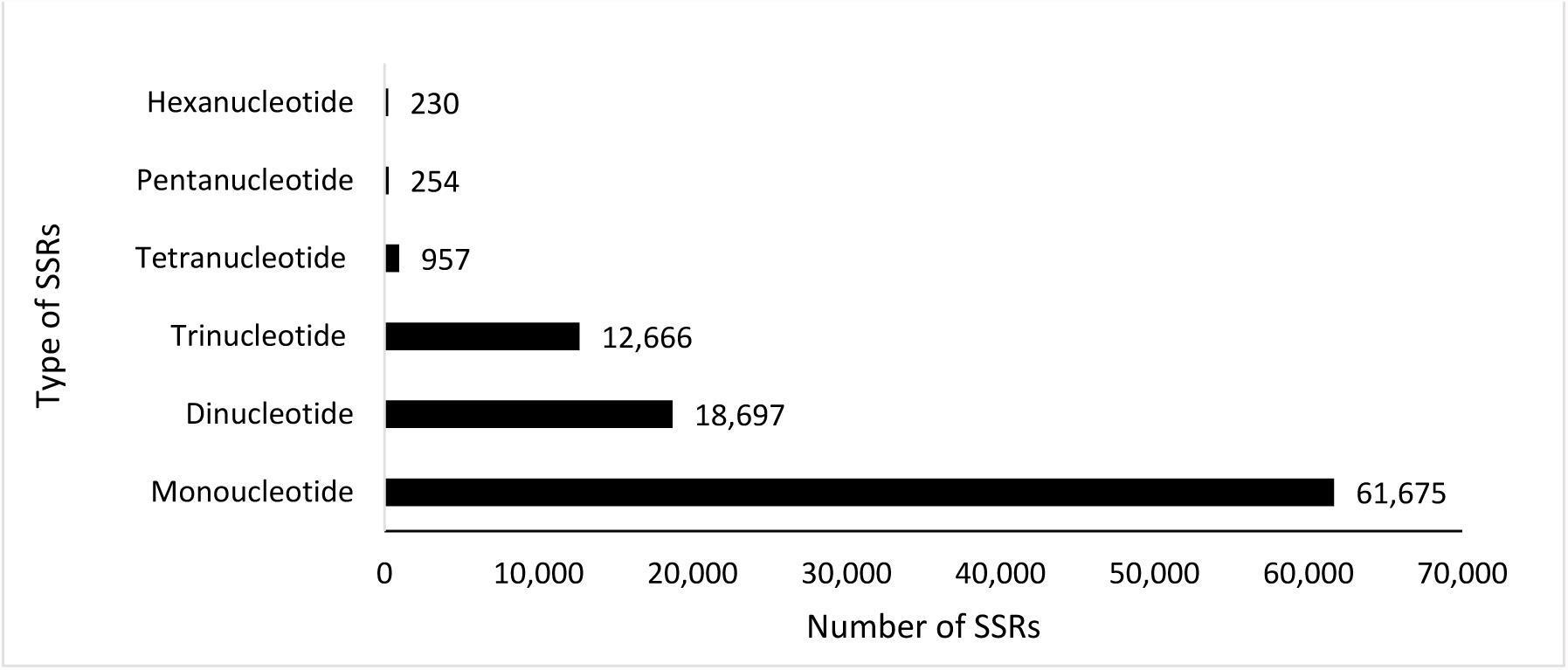
Simple sequence repeats (SSRs) classes identified in the leaf transcripts of *A. marmelos*.

SSRs are codominant markers and are well dispersed throughout plant genomes. SSRs are popularly used for marker-assisted selection, fingerprinting, diversity assessment and QTL identification [46]. Routinely, because of massively parallel sequencing, a large number of SSRs are identified in the medicinal plants via transcriptome assemblies because they are more robust and can also be transferred among different species within the same genus These SSRs identified for *A. marmelos* can be used for the fingerprinting of different accessions otherwise until recently studies were conducted in *A. marmelos* using RAPDs and ISSRs which are general markers and are not based on previous genomic information [1,47].

### 3.5. Transcriptional factors identification

Gene expression patterns are regulated by transcription factors which in turn determine the different biological process[48]. A total of 7,002 transcription factors were retrieved PlantTFDB. The fifty-two transcription factors were unique to the *A. marmelos* leaves. Although out of total 6122 were extant above hundred in the unigenes. The most abundant were Auxin response factors (ARFs) (717), Myeloblastosis (MYB-related) (562), a basic domain/leucine zipper (bZIP) (437) and basic helix–loop–helix (bHLH) (417), whereas HB-Other (132) and CAMATA (109) were least abundant (Figure 4).

**Figure 4.**
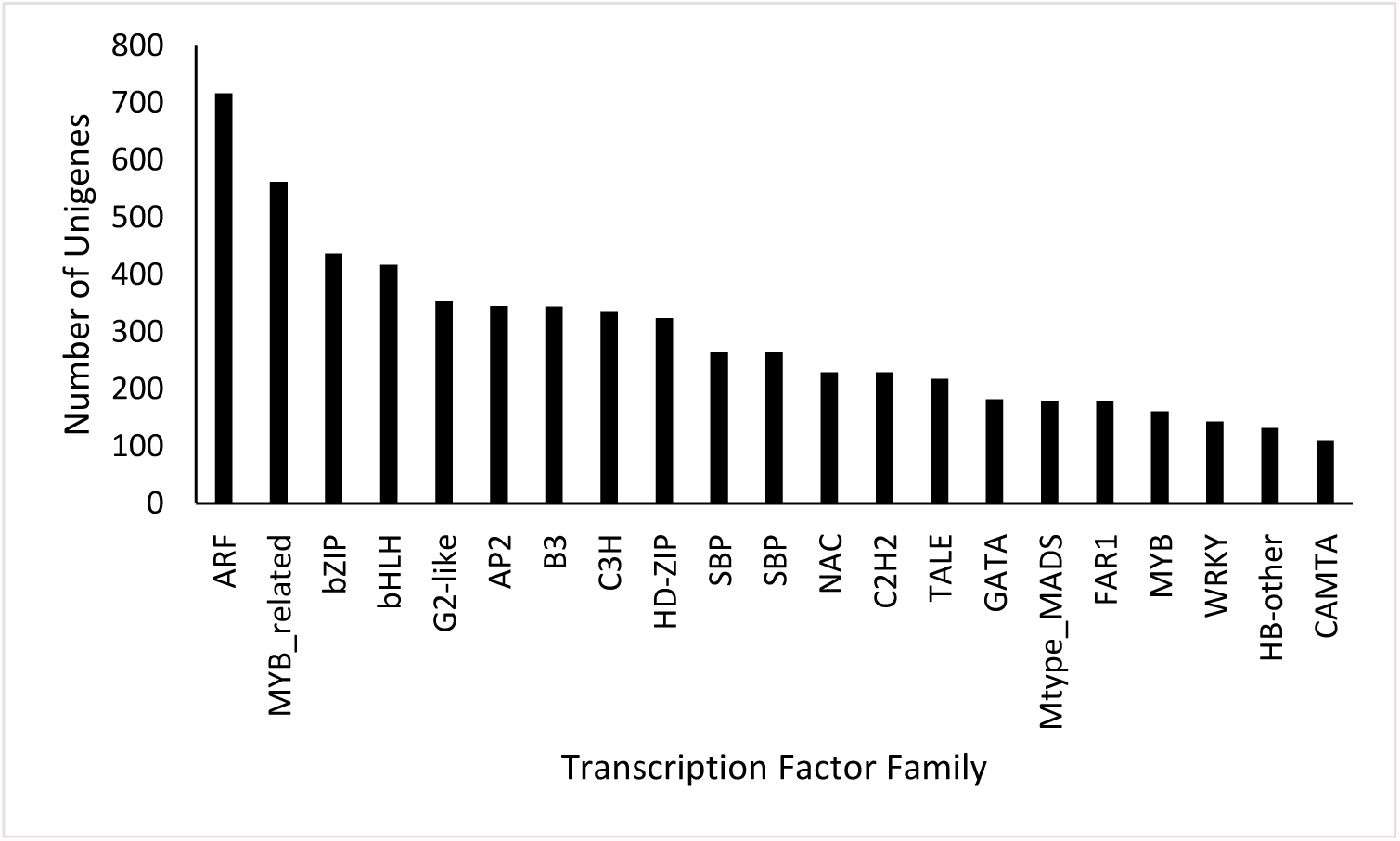
Top 21 families of transcription factors identified in the *A. marmelos* leaf.

Auxin is the plant hormone that regulates different plant process from growth to senescence. Auxin response factors are necessary for the plant to response to auxin stimuli; they channelize the response via auxin response DNA elements present in the primary auxin response genes. ARFs switch on and off the auxin response gene via their transcriptional activation domain or transcriptional repression domain[49,50]. MYB-related transcription factors play many roles in plants from plant defence including protection against biotic and abiotic stresses. MYB transcription factors also regulate the metabolism of the phenylpropanoid pathway and are well studied with respect to the regulation of primary and secondary metabolism in the plant [51,52]. Likewise, bZIP and bHLH transcription factors are also involved in the metabolic biosynthesis in plants especially by activation of phenylpropanoid genes [53,54].

### 3.6. Transcripts encoding Cytochrome p450s (CYPs) and Glucosyltransferase (GTs)

CYPs help in primary and secondary metabolism of plants by catalysing monooxygenation reactions. These cytochromes assist in the diversification of metabolic pathways in plants. Currently, these are potential targets for metabolic engineering for the overproduction of metabolites of interest [55]. There were 477 transcripts in total that were annotated a cytochrome p450s. Considering their vital role in metabolic pathway, we further analyzed the abundance of SSRs annotated within these cytochrome p450 transcripts (Table 4). Among the 128 identified SSRs, 85 were with mono repeats, 7 were with di repeats, and 36 were with tri repeats.

**Table 4.** Prediction of simple sequence repeats (SSRs) for the annotated transcripts with Cytochrome p450s (CYPs) and Glucosyltransferase (GTs).

| Description | (CYPs) | (GTs) |
| --- | --- | --- |
| Total number of annotated transcripts encoding | 477 | 314 |
| Total size of annotated transcripts (bp) | 1,111,258 | 778,664 |
| Number of SSRs identified | 128 | 247 |
| Number of annotated transcripts containing SSR | 94 | 165 |
| Number of annotated transcripts containing more than 1 SSR | 30 | 64 |
| Number of SSRs present in the compound formation | 5 | 16 |

The last step in the production of plant secondary metabolites is glycosylation which is carried out by glycotransferases (GTs) [56–58]. A total of 314 transcripts were annotated as glucosyltransferase. We analyzed the abundance of SSRs presence in these transcripts, and among the 247 identified, 109 were mono repeats, 79 were di repeats, 58 were tri repeats and only 1 was identified as a tetra repeats (Table 4). The SSRs identified using CYPs and GTs can be of immense potential for identifying genetic diversity among different *A. marmelos* accessions with divergent metabolic profiles.

### 3.7. Identification of biosynthetic pathways in A. marmelos leaf

*A. marmelos* leaves are used for the treatment of several medical conditions in Ayurveda and Yunani medicine systems [59]. The transcripts with the highest Fragments per Kilobase per Million mapped reads (FPKM) values were extracted from annotation file along with Kyoto encyclopedia of genes and genomes (KEGG) ID and sorted from the RNA-Seq by Expectation Maximization (RSEM) file that was obtained from assembly, for transcript quantification. Using the KEGG ID, pathways were identified. RSEM is commonly used to obtain information regarding transcript abundance from RNA-Seq data of an organism even without a reference genome [60]. The pathway analysis identified that monoterpenoid biosynthesis and thiamine pathways were the two most expressed pathways present in the *A. marmelos* leaves (Figure 5).

**Figure 5.**
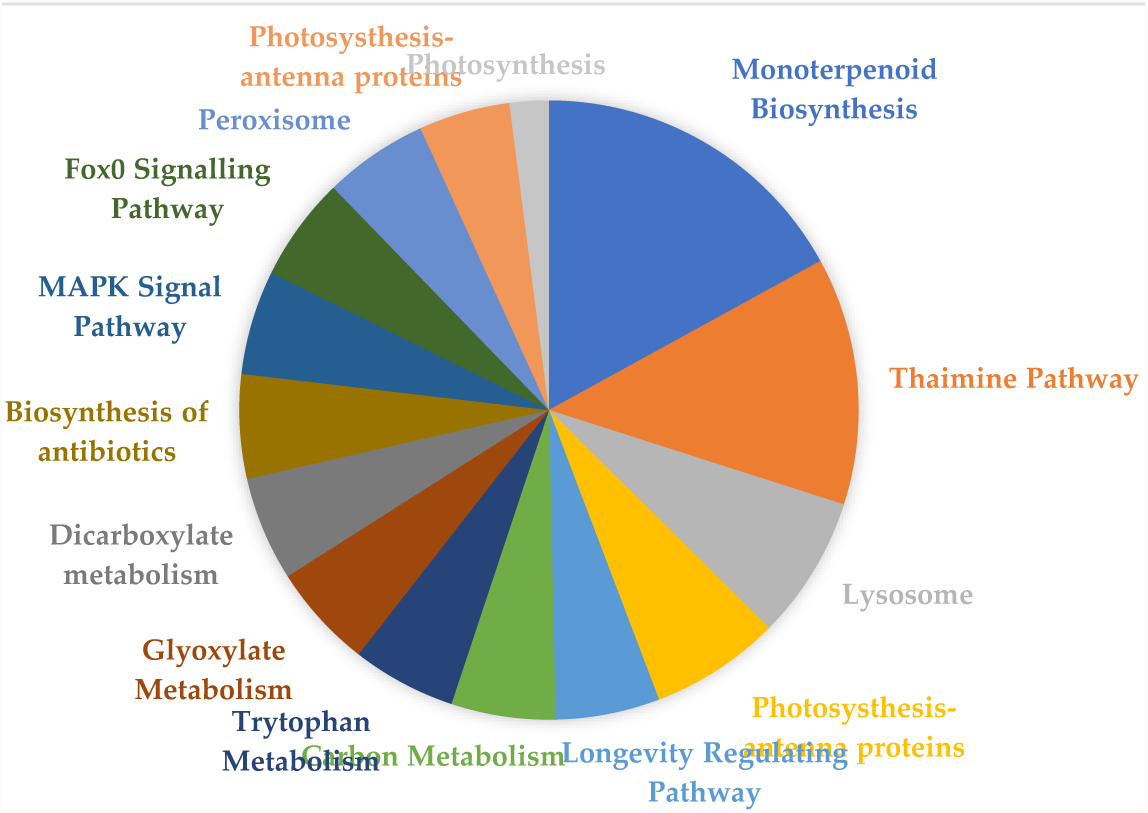
Represents the top 15 pathways in the *A. marmelos* leaf.

*A. marmelos* leaves have been reported to contain monoterpenoids as the principal metabolites in the leaves - as high as 93.9 % [61,62]. Moreover, this monoterpenoid content is not affected by the geographic location of the plant as they remain unaffected by changes in altitude, unlike many other metabolites [63]. Thiamine is naturally produced in the plants as a sulphur comprising and water-soluble compound. *A. marmelos* contains thiamine although thiamine concentration is higher in fruits than leaves and is among the fruits with highest thiamine content [64,65].

## 4. Conclusions

In case of underexploited plant species, there is not enough genomic information available to proceed for their genetic improvement and to transfer important genes from them to the cultivated crops Transcriptome assembly is a cost-effective alternative to genome sequencing for obtaining information of expressed genes and to assist in more effective development of underexploited crops and medicinal plants. RNA-seq shines light on genes and their functions, pathways present, and can subsequently lead to evolutionary studies via molecular markers. We have successfully performed and generated data of the first *de novo* transcriptome assembly of *A. marmelos*, a plant with religious, medicinal and horticultural importance [66]. It is the first-ever information about this plant which will be of immense value for evolutionary studies and development of the valuable resource for *A. marmelos*. Also, once transcriptome reference is available, anchored based transcriptome assemblies and different type of evolutionary studies can be performed within family Rutaceae involving genus *Aegle*.

